# Integrated intra- and intercellular signaling knowledge for multicellular omics analysis

**DOI:** 10.1101/2020.08.03.221242

**Authors:** Dénes Türei, Alberto Valdeolivas, Lejla Gul, Nicolàs Palacio-Escat, Olga Ivanova, Attila Gábor, Dezső Módos, Tamás Korcsmáros, Julio Saez-Rodriguez

## Abstract

Molecular knowledge of biological processes is a cornerstone in the analysis of omics data. Applied to single-cell data, such analyses can provide mechanistic insights into individual cells and their interactions. However, knowledge of intercellular communication is scarce, scattered across different resources, and not linked to intracellular processes. To address this gap, we combined over 100 resources in a single database. It covers the interactions and roles of proteins in inter- and intracellular signal transduction, as well as transcriptional and post-transcriptional regulation. We also provide a comprehensive collection of protein complexes and rich annotations on the properties of proteins, including function, localization, and role in diseases. The resource is available for human, and via homology translation for mouse and rat. The data is accessible via *OmniPath*’s web service, a Cytoscape plugin, and packages in R/Bioconductor and Python, providing convenient access options for both computational and experimental scientists. Our resource provides a single access point to knowledge spanning intra- and intercellular processes for data analysis, as we demonstrate in applications to study SARS-CoV-2 infection and ulcerative colitis.

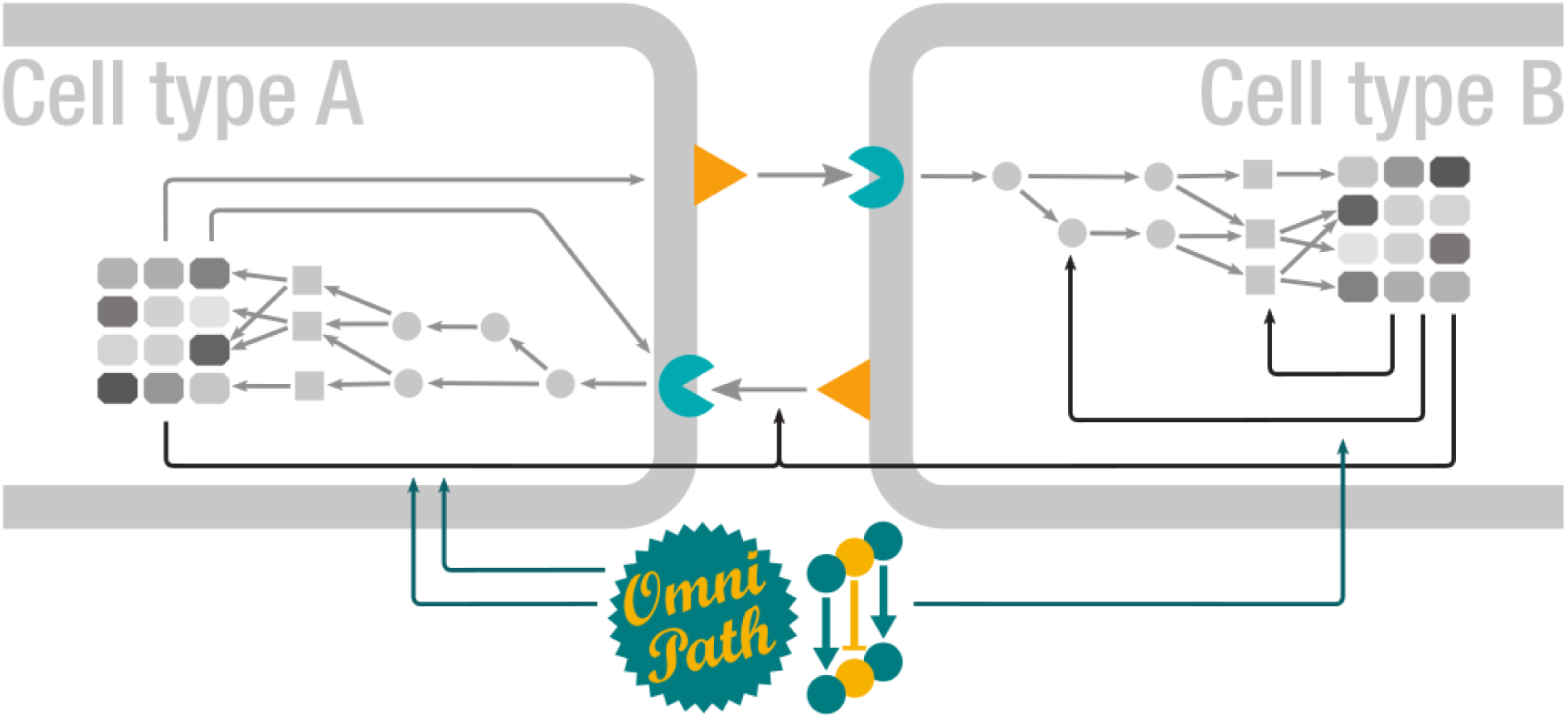

## Introduction

Cells process information by physical interactions of molecules. These interactions are organized into an ensemble of signaling pathways that are often represented as a network. This network determines the response of the cell under different physiological and disease conditions. In multicellular organisms, the behaviour of each cell is regulated by higher levels of organization: the tissue and, ultimately, the organism. In tissues, multiple cells communicate to coordinate their behavior to maintain homeostasis. For example, cells may produce and sense extracellular matrix (ECM), and release enzymes acting on the ECM as well as ligands. These ligands are detected by receptors in the same or different cells, that in turn trigger intracellular pathways that control other processes, including the production of ligands and the physical binding to other cells. The totality of these processes mediates the intercellular communication in tissues. Thus, to understand physiological and pathological processes at the tissue level, we need to consider both the signaling pathways within each cell type as well as the communication between them.

Since the end of the nineties, databases have been collecting information about signaling pathways^1^. These databases provide a unified source of information in formats that users can browse, retrieve and process. Signaling databases have become essential tools in systems biology and to analyze omics data. A few resources provide ligand-receptor interactions^2–6^. However, their coverage is limited, they do not include key players of intercellular communication such as matrix proteins or extracellular enzymes, and they are not integrated with intracellular processes. This is increasingly important as new techniques allow us to measure data from single cells, enabling the analysis of inter- and intracellular signaling. For example, the recent *CellPhoneDB*^6^ and *ICELLNET*^7^ tools provide computational methods to prioritize the most likely intercellular connections from single cell transcriptomics data, and *NicheNet*^8^ expands this to intracellular gene regulation. A comprehensive resource of inter- and intracellular signaling knowledge would enhance and expedite these analyses.

To effectively study multicellular communication, a resource should: (a) classify proteins by their roles in intercellular communication, (b) connect them by interactions from the widest possible range of resources and (c) integrate all this information in a transparent and customizable way, where the users can select the resources to evaluate their quality and features, and adapt them to their context and analyses. Prompted by the lack of comprehensive efforts addressing principle (a), we built a database on top of *OmniPath*^9^, a resource which has already shown the benefits of principles (b) and (c). This new resource focuses on intercellular communication and its integration with intracellular signaling, providing prior knowledge for modeling and analysis methods. It combines 103 resources to build on an integrated database of molecular interactions, enzyme-PTM *(post-translational modification)* relationships, protein complexes and annotations about intercellular communication and other functional attributes of proteins.

We demonstrate with two case studies that we provide a versatile resource for the analysis of single-cell and bulk omics data. First, we studied the potential influence of ligands secreted in severe acute respiratory syndrome coronavirus 2 *(SARS-CoV-2)* infection on the inflammatory response through autocrine signaling. We identified signaling mechanisms that may lead to the dysregulated inflammatory and immune response shown in severe cases. Second, we examined the rewiring of cellular communication in *ulcerative colitis* (UC) based on single cell data from the colon. By analyzing downstream signalling from the intercellular interactions, we found pathways associated with the regulatory T cells targeted by myofibroblasts in UC.

## Results

We used four major types of resources: (1) molecular *interactions*, (2) *enzyme-PTM* relationships, (3) protein *complexes* and (4) molecule *annotations* about function, localization and other attributes (Figure 1a). The *pypath* Python package combined the resources from those four types to build four corresponding integrated databases. Using the *annotations, pypath* compiled a fifth database about the roles in intercellular communication (*intercell*; Figure 1b). The ensemble of these five databases is what we call *OmniPath*, combining data from 103 resources (Figure 1a and Supplementary Table S1).

**Figure 1:**
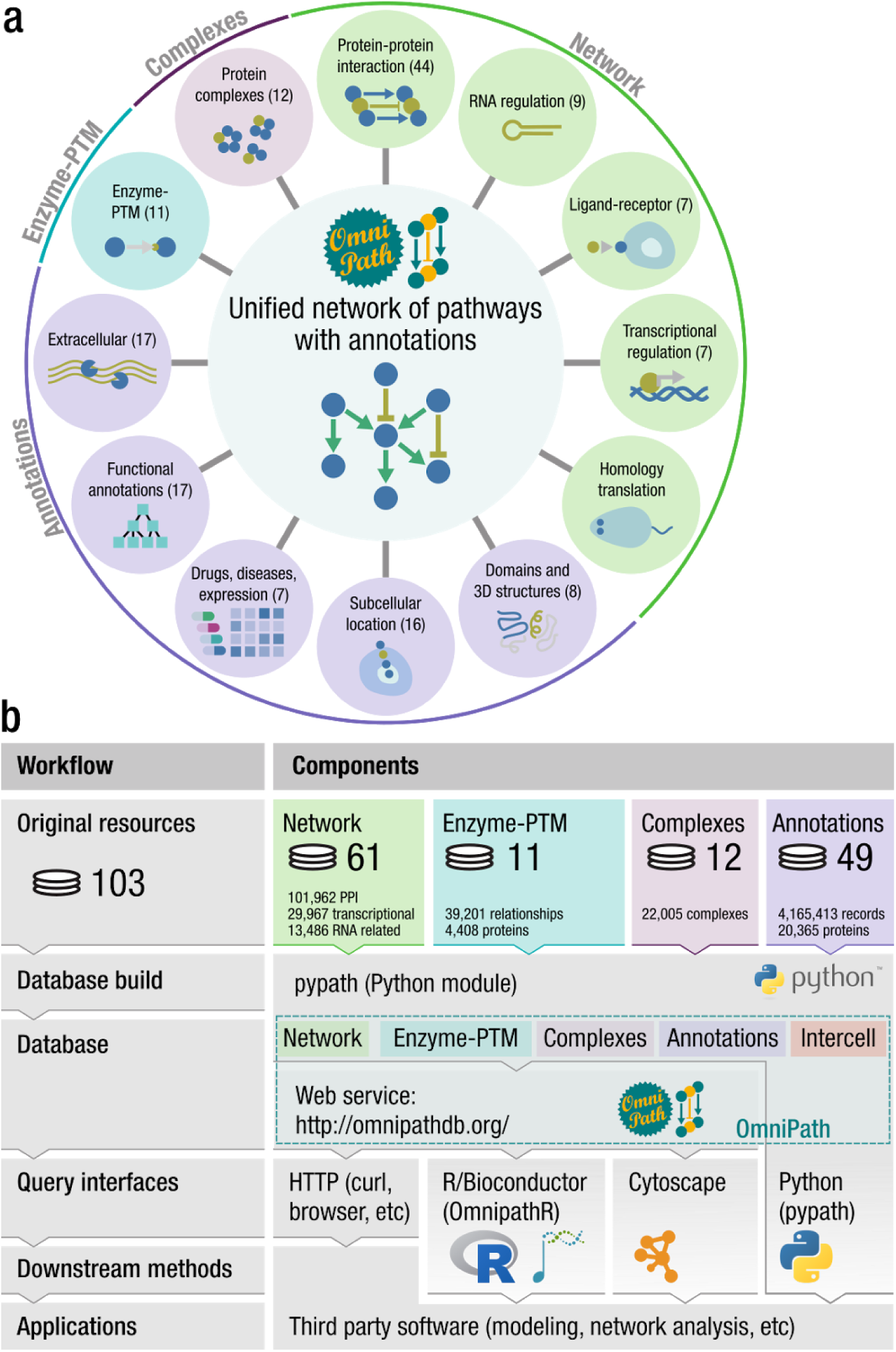
The composition and workflow of OmniPath. **(a)** Database contents with the respective number of resources in parentheses. **(b)** Workflow and design: OmniPath is based on four major types of resources, and the pypath Python package combines the resources to build five databases. The databases are available by pypath, the web resource at <u>omnipathdb.org</u>, the R package OmnipathR, the Cytoscape plug-in and can be exported to formats such as Biological Expression Language (BEL).

### A focus on intercellular signaling

To create a database of intercellular communication, we defined the roles that proteins play in this process. Ligands and receptors are main players of intercellular communication. Many other kinds of molecules have a great impact on the behavior of the cells, such as matrix proteins and transporters. We defined eight major (Figure 2) and 17 minor generic functional categories of intercellular signaling roles (Supplementary Table S6 and S10). We also defined ten locational categories (e.g. *plasma membrane peripheral*), using in addition structural resources and prediction methods to annotate the transmembrane, secreted and peripheral membrane proteins. Furthermore, we provide 994 specific categories (e.g. *neurotrophin receptors*). Each generic category can be accessed by resource (e.g. *ligands from HGNC*) or as the combination of all contributing resources (Supplementary Figure S5). To provide highly curated annotations, we checked every entry in each category manually against UniProt datasheets to exclude wrong annotations. Overall we defined 1,170 categories and provided 54,330 functional annotations about intercellular communication roles of 5,781 proteins.

**Figure 2:**
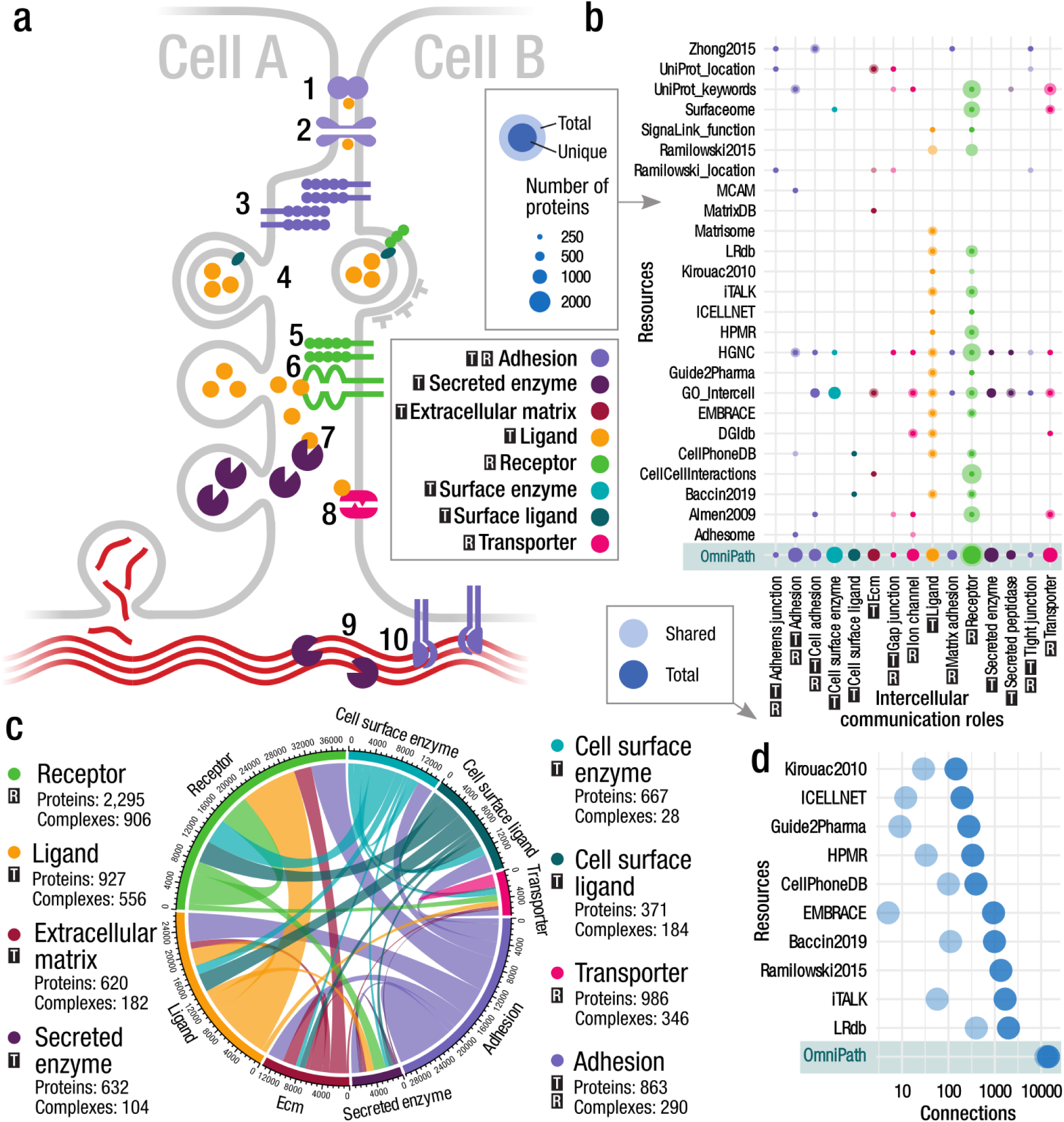
The composition and representation of the intercellular signaling network. We assigned intercellular communication roles to proteins based on evidence from multiple resources. In all panels: T -transmitter; R -receiver. **(a)** Schematic illustration of the intercellular communication roles and their possible connections. Cells are physically connected by proteins forming tight junctions (1), gap junctions (2) and other adhesion proteins (3); they release vesicles which can be taken up by other cells (4); some receptors form complexes (5) to detect secreted ligands (6); transporters might also be affected by factors released by other cells (8); enzymes released into the extracellular space act on ligands and the extracellular matrix (7, 9); cells release the components of the extracellular matrix and bind to the matrix by adhesion proteins (10). **(b)** The main intercellular communication roles (x axis) and the major contributing resources (y axis). Size of the dots represents the number of proteins annotated to have a certain role in a given resource. The darker areas represent the overlaps (proteins annotated in more than one resource for the same role) while the lighter color denotes those unique to that resource. **(c)** The intercellular communication network. The circle segments represent the eight main intercellular communication roles. The edges are proportional to the number of interactions in the OmniPath PPI network connecting proteins of one role to the other. **(d)** Number of unique, directed transmitter-receiver (e.g. ligand-receptor) connections by resources.

We collected the proteins for each intercellular communication functional category using data from 27 resources (Supplementary Table S6). Combining them with molecular interaction networks from 48 resources (Supplementary Table S2) we created a corpus of putative intercellular communication pathways (Figure 2c). To have a high coverage on the intercellular molecular interactions, we also included ten resources focusing on ligand-receptor interactions (Figure 3, Supplementary Figure S1).

**Figure 3:**
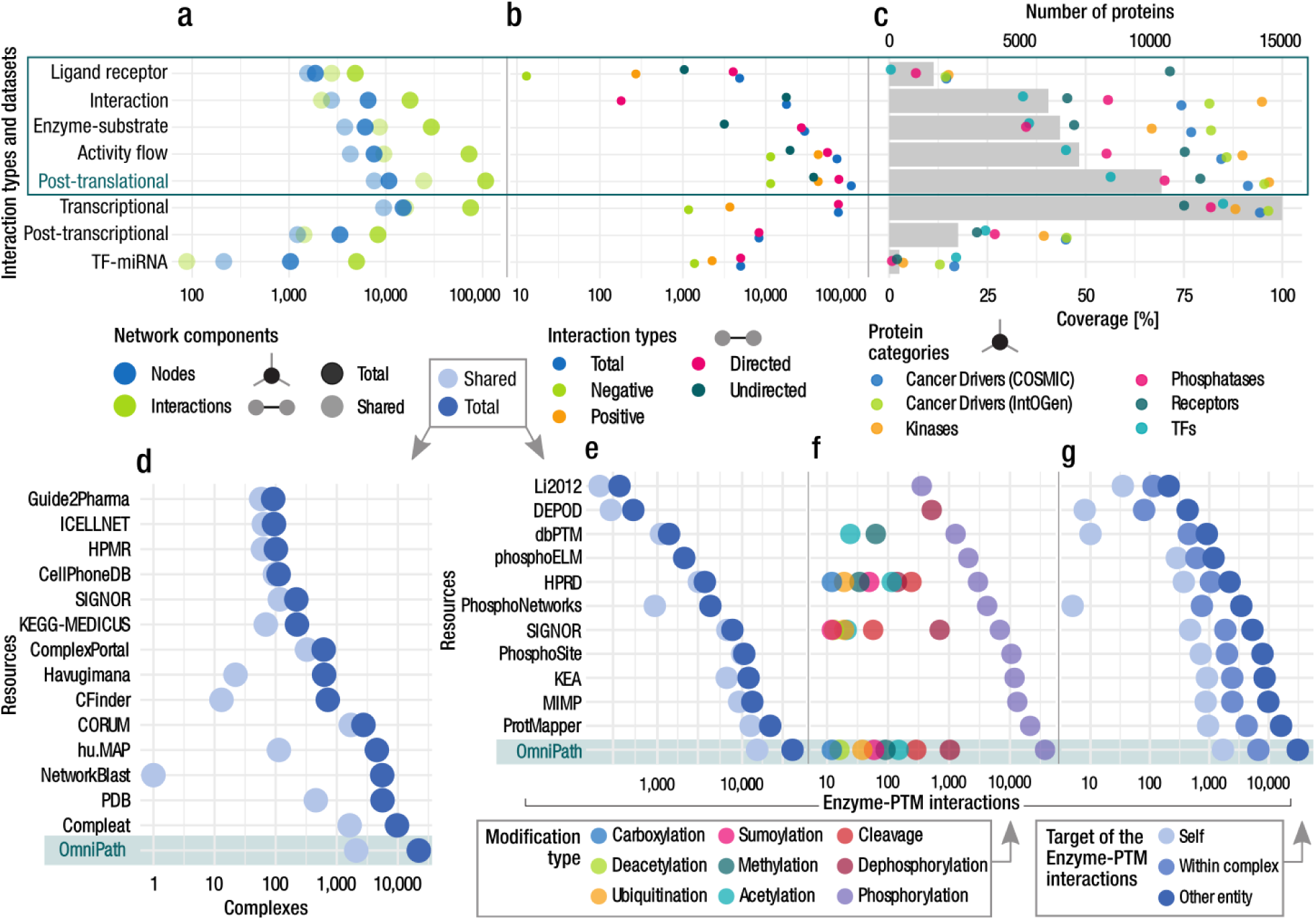
Quantitative description of the network, complex and enzyme-PTM databases. **(a-c)** Networks by interaction types and the network datasets within the PPI network. **(a)** Number of nodes and interactions. The light dots represent the shared nodes and edges (in more than one resource), while the dark ones show their total numbers. **(b)** Causality: number of connections by direction and effect sign. **(c)** Coverage of the networks on various groups of proteins. Dots show the percentage of proteins covered by network resources for the following groups: cancer driver genes from COSMIC and IntOGen, kinases from kinase.com, phosphatases from Phosphatome.net, receptors from the Human Plasma Membrane Receptome (HPMR) and transcription factors from the TF census. Gray bars show the number of proteins in the networks. The information for individual resources is in Supplementary Figures S1-3. **(d-g)** On each panel the bottom rows represent the combined complex and enzyme-PTM databases contained in OmniPath **(d-e)**. Number of complexes **(d)** and enzyme-PTM **(e)** interactions by resource. **(f)** Enzyme-PTM relationships by PTM type. **(g)** Enzyme-PTM interactions by their target. Light, medium and dark dots represent the number of enzyme-PTM relationships targeting the enzyme itself, another protein within the same molecular complex or an independent protein, respectively.

Many of the proteins in intercellular communication work as parts of complexes. We therefore built a comprehensive database of protein complexes and inferred their intercellular communication roles: a complex belongs to a category if and only if all members of the complex belong to it. We obtained 14,348 unique, directed transmitter-receiver (e.g. ligand-receptor) connections, around seven times more than the largest of the resources providing such kind of data (Figure 2d). This large coverage is achieved by not only integrating ten ligand-receptor resources, but also complementing these with data from annotation and interaction resources.

An essential feature of this novel resource is that it combines knowledge about intercellular and intracellular signaling. Thus, using *OmniPath* one can, for example, easily analyze the intracellular pathways triggered by a given ligand or check the transcription factors (TFs) and microRNAs (miRNAs) regulating the expression of such ligands.

#### *OmniPath*: an ensemble of five databases

The abovementioned intercellular database exists in *OmniPath* together with four further databases (Figure 1b), supporting an integrated analysis of inter- and intracellular signaling.

##### The network of molecular interactions

The *network* database part covers four major domains of molecular signaling: (i) protein-protein interactions (PPI), (ii) transcriptional regulation of protein-coding genes, (iii) miRNA-mRNA interactions and (iv) transcriptional regulation of miRNA genes (TF-miRNA). We further differentiated the PPI data into four subsets based on the interaction mechanisms and the types of supporting evidence: 1) literature curated activity flow (directed and signed; corresponds to the original release of *OmniPath*^9^), 2) activity-flow with no literature references, 3) enzyme-PTM and 4) ligand-receptor interactions (Figure 3a-c). In total, the resource contained 103,396 PPI interactions between 12,469 proteins from 38 original resources (Supplementary Table S2). The large number of unique interactions added by each resource underscores the importance of their integration (Supplementary Figures S2-4). The interactions with effect signs, essential for mechanistic modeling, are provided by the activity flow resources (Figure 3b). The combined PPI network covered 53% of the human proteome (SwissProt), with an enrichment of kinases and cancer driver genes (Figure 3c). The transcriptional regulation data in *OmniPath* was obtained from *DoRothEA*^10^, a comprehensive resource of TF regulons integrating data from 18 sources. In addition, six literature curated resources were directly integrated into *OmniPath* (Supplementary Table S8). The miRNA-mRNA and TF-miRNA interactions were integrated from five and two literature curated resources, with 6,213 and 1,803 interactions, respectively. Overall, we included 61 network resources in *OmniPath* (Supplementary Table S2).

Furthermore, *pypath* provides access to additional resources, including the Human Reference Interactome^11^, ConsensusPathDB^12^, Reactome^13^, ACSN^14^ and WikiPathways^15^.

##### Enzyme-PTM relationships

In enzyme-PTM relationships, enzymes (e.g. kinases) alter specific residues of their substrates, producing so-called post-translational modifications (PTM). We combined 11 resources of enzyme-PTM relationships mostly covering phosphorylation (94% of all) and dephosphorylations (3%) (Figure 3f). Overall, we included 39,201 enzyme-PTM relationships, 1,821 enzymes targeting 16,467 PTM sites (Figure 3e-g). Besides phosphorylation and dephosphorylation, only proteolytic cleavage and acetylation account for more than one hundred interactions. Most of the databases curated only phosphorylation, and *DEPOD*^16^ exclusively dephosphorylation. Only *SIGNOR*^17^ and *HPRD*^18^ contained a large number of other modifications (Figure 3f). 60% of the interactions were described by only one resource, and 92% of them by only one literature reference (Figure 3e). Self-modifications, e.g. autophosphorylation and modifications between members of the same complex comprised 4% and 18% of the interactions, respectively (Figure 3g).

##### Protein complexes

The *complexes* database of *OmniPath* included 22,005 protein complexes described by 12 resources from 4,077 articles (Figure 3d). A complex is defined by its unique combination of members. 14% of them were homomultimers, 54% had four or less unique components while 20% of them had 18 or more. 71% of the complexes had stoichiometry information.

##### Annotations: function, structure and localization

Annotations provide information about the function, structure, localization, classification and other properties of molecules. We compiled the *annotations* database from 49 resources. The format of the records from each of these resources is different. The simplest ones only define a category of proteins, like *Cell Surface Protein Atlas*^19^ *(CSPA)* that collects the proteins localized on the cell surface. More complex annotation records express a combination of multiple attributes. For example, each of the annotations from the *Cancer Pathway Association Database*^20^ *(CPAD)* contain seven attributes to describe a relationship between a protein or miRNA, a pathway and their effect on a specific cancer type (Supplementary Figure S4). The pathway and gene sets are also part of the annotation database, as these are useful for functional characterization of omics data and enrichment analysis.

Overall, the *annotations* database included 5,475,532 records about 20,365 proteins, virtually the whole protein-coding genome, 19,566 complexes and 182 miRNAs. The majority of the annotations for complexes were the result of our *in silico* inference: if all members of a complex share a certain annotation we assign this annotation to the complex itself.

The *annotations* database can be used in different ways: selecting one resource, its data can be reconstituted into a conventional data frame with attributes as columns and annotations as rows. Alternatively, specific sets of proteins can be queried e.g. “the members of the *Notch pathway* according to *SIGNOR*”^17^ or “the *hypoxia upregulated* genes according to *MSigDB*”^21^.

##### Homology translation to rodents

*OmniPath* comprises human resources. We translated the network and the enzyme-PTM relationships to mouse and rat by protein homology using *NCBI HomoloGene*, covering 81% and 31% of the interactions for mouse and rat, respectively (Supplementary Table S9). In addition, *pypath* is able to translate to other organisms.

### Case Studies

*OmniPath* provides a single-access point to resources covering diverse types of knowledge. Thus, it can be used as an input to many analysis tools, and is particularly useful for tools that span over molecular processes typically considered separately (Figure 4a). To illustrate this, we used two examples where we extracted from *OmniPath* different types of intra- and intercellular knowledge for computational analysis of bulk and single-cell RNA-Seq data.

**Figure 4:**
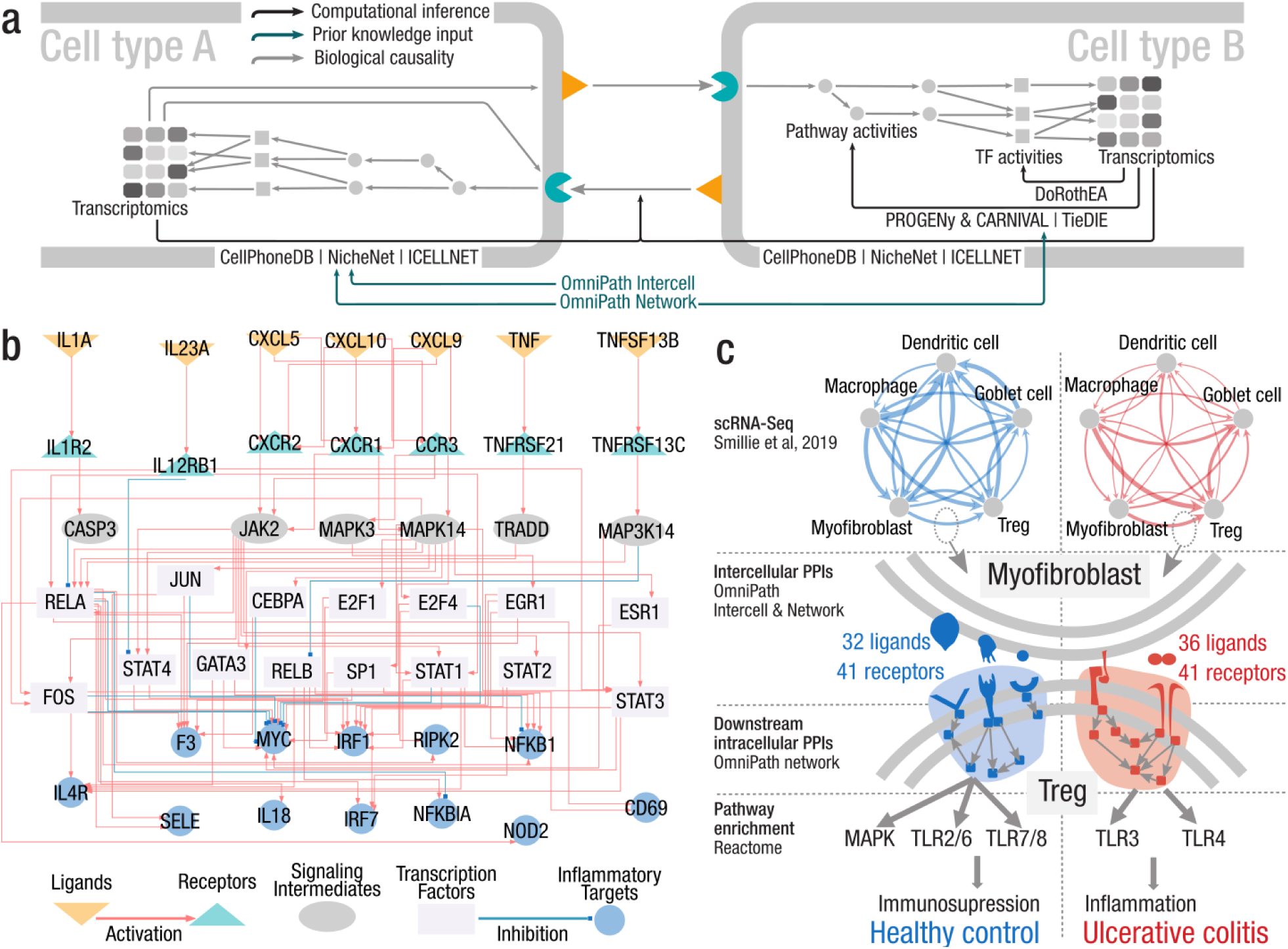
Illustrations of the integrated analysis of inter- and intracellular signaling. **(a)** Examples of tools for integrated analysis of tissue level signaling from cell type specific transcriptomics data that can be applied with the prior knowledge available in OmniPath. **(b)** Inter and intracellular signaling interactions linking the top predicted ligands over-expressed after SARS-CoV-2 infection to their potential immune response targets in the Calu3 cell line. Top-ranked ligands (orange) connect to their potential receptors (turquoise) that trigger an intracellular cascade until reaching TFs (light grey), that in turn regulate the expression of the target genes (blue). Signaling intermediates (dark grey) connect receptors to TFs across their shortest path. **(c)** Intercellular connections and their downstream effect in UC compared to healthy control. Top: communication network of five cell types reconstructed from scRNA-Seq; the thickness of the edges is proportional to the number of intercellular connections. Bottom: condition specific ligand-receptor connections between myofibroblasts and regulatory T cells trigger a immunosuppressive versus an inflammatory signaling in T cells, in healthy and UC, respectively.

#### Analysis of intra- and intercellular processes in SARS-CoV-2 infected lung epithelial cancer cells

*NicheNet* is a recently developed method to prioritize ligand-target relationships between interacting cells by combining their expression data with prior knowledge on interaction networks^8^. For this purpose, *NicheNet* explores the most consistent inter- and intracellular protein interactions in accordance with a given gene expression dataset. In the *NicheNet* publication, the authors collected different types of interactions from more than 20 databases to build a ligand-receptor network, a signaling network and a gene regulatory network. Here, we built a network for analysis with *NicheNet* using exclusively *OmniPath*.

We used this network to investigate the mechanistic processes leading to the excessive inflammatory innate response and dysregulated adaptive host immune defense that may occur in severe *COVID-19* cases^22^. We studied the autocrine regulatory effect of ligands secreted in *SARS-CoV-2* infection of epithelial lung cancer cells *(Calu3)* on the expression of inflammatory response genes (Methods and Supplementary Note 1, data from^23^). Out of a total of 117 ligands over-expressed in *SARS-CoV-2* infection according to *NicheNet*, we selected the 12 top-ranked ones for subsequent analysis (Supplementary Figure 6). Among them, we found various cytokines: interleukins (*IL23A* and *IL1A*), tumor necrosis factors (*TNF* and *TNFSF13B*) and chemokines (*CXCL5*, *CXCL9* and *CXCL10*), known to be involved in the inflammatory response. The top predicted target genes for these 12 ligands were enriched for inflammatory response gene sets (average p-value=3.25e-08 from Fisher’s exact tests after 10 cross-validation rounds). Then, we explored the signaling events linking these ligands to their target genes (Figure 4b, Methods and Supplementary Note 1). We identified several key proteins of the *JAK-STAT pathway*, a main regulator of the inflammatory response, that has been suggested as a potential target to treat *COVID-19*^24^. We also found ligands that potentially trigger the *MAPK pathway* that has also been reported to be promoted by *SARS-CoV-2* infection^25, 26^. We found further support for these results in the literature (Supplementary Note 1).

#### Alteration of intercellular communication in ulcerative colitis

As a second case study, we used single-cell RNA-Seq data ^27^ from *ulcerative colitis (UC)* to investigate paracrine signaling using *OmniPath.* We explored the intercellular interactions comparing the healthy state and non-inflamed UC. We selected five interacting cell types: dendritic cell, macrophage, regulatory T cell (Treg), myofibroblast and Goblet cell. Combining the expression data with *OmniPath*, we built a network of communication between these five cell types and quantified the disease specific changes. Then, we added the components from the *OmniPath* PPI network two steps downstream of the cell type specifically expressed receptors. Finally, we performed a pathway enrichment analysis using *Reactome*^13^; Methods).

We found that in healthy condition dendritic cells (DC) were tightly connected to the four other cell types. In contrast, in UC the connections shifted towards the Treg cells instead of DC, in agreement with previous findings^27^ (Figure 4c). We found a 30% increase in the amount of ligand-receptor and ligand-adhesion interactions between myofibroblasts and Treg in UC versus healthy, even though the number of connections is similar in both conditions. In an analysis of downstream signaling in Treg cells we found pathways known to downregulate the pro-inflammatory function of Treg cells to be active in healthy state, including the *MAPK*^28^, *TLR2*^29^ and *TLR7*^30^ *pathways* (Supplementary Table S11). In contrast, the pro-inflammatory *TLR4*^31^ and *TLR3 pathways*^32^ were upregulated in UC. These results suggest a pro-inflammatory response in UC, where the anti-inflammatory role of regulatory T cells is deteriorated by myofibroblasts.

## Discussion

### A single access point to annotated causal knowledge

Here, we present a single-access point to over 100 resources containing prior knowledge of intra- and intercellular processes, building on the *OmniPath* framework. To achieve this, we developed versatile annotations of intercellular communication roles, combined with a network covering intra- and intercellular signaling as well as gene regulation.

We focused on resources that follow the *activity flow* representation where nodes are linked with signed and directed edges representing a certain influence. The alternative *process description* representation describes the underlying processes as biochemical reactions^33^. Integrative resources such as *STRING*^34^, *PathwayCommons*^35^, *ConsensusPathDB*^12^, *PathMe* and *ComPath*^36^ use mostly the major process description resources (e.g. *Reactome*^13^ and *ACSN*^14^) and resources with undirected interactions (e.g. *IntAct*^37^ and *BioGRID*^38^), and only few activity flow resources.

However, for many applications, *process description* representation must undergo a conversion to *activity flow* representation^35^. This conversion is technically challenging, leads to information loss^35, 39, 40^, alters the network topology, and affects downstream applications. On the other extreme, undirected interactions lack information about directionality and stimulatory and inhibitory effects, which are essential for many analytical methods, in particular those that aim to capture causal relationships. The activity flow representation is between both: interactions are presented as signed and directed edges, regardless of the underlying biochemistry. Due to this abstraction, activity flow has limitations and the stimulatory and inhibitory nature of the interactions can be ambiguous^41^. Despite these limitations, activity flow databases are widely used because their level of abstraction provides a convenient input for multiple analysis techniques^42^.

#### Comprehensive knowledge for multicellular omics analysis

As we demonstrated here, *OmniPath* is able to provide the input knowledge for different data analysis tools, such as *CellPhoneDB*^6^*, NicheNet*^8^ and *CARNIVAL*^43^ to infer communication between and within cell types. *OmniPath* is not limited to literature curated interactions and it includes activity flow, kinase-substrate and ligand-receptor interactions without references as separate datasets, so that the users can decide which ones to use according to their purposes.

The rich annotations allow users to dive into specific knowledge and extract information across resources. Information obtained via text-mining approaches^44, 45^ can be used to complement the curated knowledge captured within *OmniPath*.

As our case studies illustrate, *OmniPath* can replace the tedious collection of information from many different databases. In the first case study, we modelled with *NicheNet* the autocrine signaling after *SARS-CoV-2* infection. Our results suggest potential signaling mechanisms leading to the dysregulated inflammatory and immune response characteristic of severe *COVID* cases. In the second study, we illustrated how conveniently *OmniPath* supports a combined analysis of inter- and intracellular signaling from single-cell transcriptomics data.

In summary, we provide a new integrated resource of biological knowledge particularly valuable for network analysis and modeling of bulk and single-cell omics data. Furthermore, with the emergence of spatially resolved omics data^46^, we anticipate that this prior knowledge of inter- and intracellular communication will be valuable to study tissue architecture.

## Methods

### Terminology

In the manuscript we use consistently the following three definitions to describe the structure of *OmniPath*:

- **database:** collection of similar records in a uniform format integrated from multiple resources (network, enzyme-PTM, complexes, annotations, intercell)
- **dataset:** a subset or variant of a database, e.g. the transcriptional interaction network is a dataset of the network database
- **resource:** any data source we use for building the databases

### Database build

To build *OmniPath* we developed a free software, the *pypath* Python module (https://github.com/saezlab/pypath, version 0.11.20). We built each segment of the database by the corresponding submodules and classes in *pypath.* In addition to the database building process, all modules rely on common utility modules from *pypath* such as the identifier translator or the downloading and caching service. *Pypath* downloads all data from the original sources. Many resources integrate data from other resources, we call these secondary resources and their relationships are listed in Supplementary Table S7.

#### Network

For the *OmniPath* network, we converted the identifiers of the different molecules and merged their pairwise connections, preserving the literature references, the information about the direction and effect sign (activation or inhibition).

In *OmniPath*, we included nine network datasets built from 61 resources (Supplementary Table S2). The first four datasets provide PPI (*‘post_translational’* in the web service) while the others transcriptional and post-transcriptional regulation. At each point below we highlight the label of the dataset in the web service.

1. We compiled the “**omnipath”** network as described in Türei et al^9^. Briefly, we combined all resources we could get access to, that are literature curated and are either activity flow, enzyme-PTM or undirected interaction resources. We also added network databases with high-throughput data. Then we added further directions and effect signs from resources without literature references.
2. The **“kinaseextra”** network contains additional kinase-substrate interactions without literature references. The direction of these interactions points from the enzyme to the substrate.
3. In the **“pathwayextra”** network, we combined further activity flow resources without literature references. However, they are manually curated and many have effect signs.
4. In the **“ligrecextra”** network, we provide additional ligand-receptor interactions from large, comprehensive collections.
5. The **“dorothea”** network comes from DoRothEA database, a comprehensive resource of transcription factor-gene promoter interactions from literature curated databases, high-throughput experiments, binding motif and gene expression-based *in silico* inference, overall 18 resources^10^. We included the interactions from DoRothEA subclassified by confidence levels from A to D, excluding the lowest confidence level E. In *OmniPath* users are able to filter the TF-target interactions by confidence level.
6. Transcriptional regulation (**“tf_target”**) directly from 6 literature curated resources. We show the size of the TF-target network at different settings in Supplementary Table S8.
7. In the **“post_transcriptional”** network, we combined 5 literature curated miRNA-mRNA interactions.
8. Transcriptional regulation of miRNA (**“tf_mirna”**) from 2 literature curated resources.
9. lncRNA-mRNA interactions from 3 literature curated resources (**“lncrna_mrna”**).

#### Enzyme-PTM interactions

After translating the identifiers, we merged enzyme-PTM interactions from 11 databases (Supplementary Table S3) based on the identity of the enzyme, the substrate and the modified residue and its position. In addition, we discarded the records where the residue could not be found in any of the isoform sequences from UniProt^47^. For each enzyme-PTM interaction, we included the original sources and the literature references. We also kept the records without literature support, e.g. from high-throughput screenings or in silico prediction.

#### Complexes

We combined 12 databases to build a comprehensive set of protein complexes (Supplementary Table S4). Seven of these databases provide information about the stoichiometry of the complexes while three contain only the lists of components. We translated the names of the components to UniProtKB accession numbers. We merged the complexes based on their identical sets of components and preserved the stoichiometry if available. We represent each complex by the UniProt IDs of their components sorted alphabetically, separated by dashes and prefixed with ‘COMPLEX:’. From the original sources, we kept the literature references, the human readable names (synonyms) and the PDB structure identifiers if available.

#### Annotations

Annotation resources provide diverse information about the localization, function or other characteristics of the molecules. We obtained annotations from 49 databases (Supplementary Table S5). For these databases, we translated IDs and extracted the fields with relevant information. Due to the heterogeneous nature of the data, in the annotation database, the content of the resources is not merged, but rather all entries are provided independently.

Each annotation record assigns one or more attributes to a molecule. One protein might have more than one annotation record from the same database. For example, Vesiclepedia provides two attributes: the vesicle type and the tissue where the protein has been detected. We combined the annotation resources into a uniform table where one column is the name of the attribute and the other is the value. As one record might have multiple attributes the records are identified by unique numbers (Supplementary Figure S4). Providing the data in this format in our web service, it can be easily reconstituted to conventional tables with standard tools like tidyr (https://tidyr.tidyverse.org) in R or pandas (https://pandas.pydata.org) in Python.

#### Complex annotations

Only four resources curate annotations of protein complexes, from these we processed the complex annotations as we did for proteins. Furthermore, we inferred annotations for complexes based on the annotations of their components. We assigned the annotations to the complex if all components agreed in all attributes that we considered relevant e.g. if all members of a complex belong to the WNT pathway then the complex is also annotated as a member of the WNT pathway.

#### Intercellular signaling roles

From the resources used in *annotations*, we selected 26 with function, location or structure information relevant in intercellular signaling. The relevant attributes we processed and combined to account for main roles in intercellular communication (e.g. ligand, receptor, ECM proteins) as well as the locational and topological properties (e.g. secreted, transmembrane). In addition, we built Boolean expressions from Gene Ontology terms to define the same categories. Overall we created 25 functional and 10 locational categories (Supplementary Table S6). Each category carries the attributes described in Supplementary Table S10 (Supplementary Figure S5). We manually checked the members of all the annotation groups, relying on literature knowledge and UniProt datasheets^47^, discarding the wrong annotations. We provide the classification of proteins and complexes by these categories in the *intercell* query of the web service.

#### Identifier translation

For each type of molecule, we chose a reference database: for proteins the UniProtKB ACs while for miRNAs the miRBase mature ACs. From these databases we obtained a reference set of identifiers for each type of molecular entity and organism. We then used translation tables provided by them to map other kinds of identifiers to the reference set. For UniProt, we corrected for deprecated or secondary ACs by translating to primary gene symbol and then to primary UniProt AC. We applied corrections to handle non-standard notations (e.g. extra dashes, greek letters). We also accounted for the ambiguity in the mapping, i.e. if one foreign identifier may correspond to multiple reference identifiers we keep all target identifiers in *OmniPath*.

#### Translation by homology to rodent species

The homology translation in *pypath* uses the NCBI HomoloGene database^48^. Because HomoloGene uses RefSeq IDs, the translation takes three steps: from UniProt to RefSeq, then to the homologous RefSeq and finally from RefSeq to UniProt. The success rate of this translation is around 80% for mouse and roughly 30% for rat (Supplementary Table S9). We translated the network data and the enzyme-PTM interactions from human to mouse and rat, the two most popular mammalian model organisms. In addition, we looked up PTMs in PhosphoSite^49^ which provides homology data for PTM sites. Then we checked the residues in the UniProt sequences^47^, and discarded the ones that did not match. The homology-translated data is included also in the *OmniPath* web service.

#### Data download and caching

At the database build we download all input data directly from the original sources (Supplementary Table S1). Certain databases either temporarily or ultimately went offline; we deposited their data in the *OmniPath Rescued Data Repository* (http://rescued.omnipathdb.org/). *Pypath* contains the URLs for all resources used including the identifier translation tables. It automatically downloads, extracts and preprocesses the data for each operation. Then it stores the downloaded data in a local cache directory which belongs to the user account on the computer. Once cache is created, *pypath* reads from it and performs the download only if requested by the user.

**Joint analysis of intra- and intercellular processes in *SARS-CoV-2* infection** The NicheNet method^8^ was built, trained and applied to a case study using interactions and annotations from *OmniPath* resources. This information was downloaded via our R package, *OmnipathR*.

#### Network construction

NicheNet generates a model based on prior knowledge to describe potential regulatory effects of ligands on target genes. To reproduce their procedure, we first built three networks accounting for protein interactions of different categories retrieved from *OmniPath*:

1. **Ligand-receptor network**: we downloaded the **“ligrecextra”** network which specifically contains known interactions between ligands and receptors. In addition, we selected proteins annotated as *ligands* or *receptors* as their main “**intercellular signaling role”**. Then we extended this network with PPI whose source is a ligand and its target a receptor.
2. **Signaling network:** we retrieved PPI from the original *OmniPath* network^9^, the **“kinaseextra”** network and the **“pathwayextra”** network.
3. **Gene regulatory network:** we selected TF-target interactions with confidence level A, B and C from the DoRothEA dataset of the **“transcriptional”** network of *OmniPath*.

Then, we computed ligand–target regulatory potential scores based on the topology of our aforementioned networks, following the protocols described in the NicheNet original study and using its associated *nichenetr* package^8^. Briefly, Personalized PageRank was applied on the union of the ligand-receptor and signaling networks considering every individual ligand as starting node. To estimate the impact of every ligand in the expression of target genes, a matrix containing the PageRank scores is multiplied by the weighted adjacency matrix of the gene regulatory network.

#### Analysis of altered ligands and pathways

We applied our OmniPath-based version of NicheNet analyses on RNA-seq data of a human lung cell line, Calu3 (GSE147507)^23^. In this study, differential expression analysis at the gene level between controls and SARS-CoV-2 infected cells was carried out using the *DESeq2* package^50^. We selected over-expressed ligands (adjusted p-value < 0.1 and Log2 fold-change > 1) after SARS-CoV-2 infection for further analysis. Then, we executed Gene Set Enrichment Analysis (GSEA) taking the Wald statistic and the hallmark gene sets from MsigDB as inputs using the *fgsea* package^51^. Inflammatory response appeared as one of the top enriched sets. We therefore selected the leading edge genes of inflammatory response, i.e. genes contributing the most to the enrichment of this particular set, as potential targets of the over-expressed ligands.

Ligand activity analysis on the aforementioned samples was conducted using the *nichenetr* package^8^. We then selected the shortest paths between receptors (the ones interacting with the top predicted ligands) and transcription factors (the ones regulating the expression of the inflammatory target genes). These paths were exported to Cytoscape^52^ to generate Figure 4b.

### Intercellular communication in ulcerative colitis

#### Intercellular interactions from OmniPath

We downloaded intercellular interactions using the ‘*import_intercell_network()*’ method in *OmnipathR* and filtered for direct cell-cell connections: we discarded extracellular matrix proteins, extracellular matrix regulators, ligand regulators, receptor regulators and matrix adhesion regulators and kept only membrane-bound (transmembrane or peripheral site of the membrane) proteins. This resulted in connections involving ligands, receptors, junction, adhesion, ion channel, transporter and cell surface or secreted enzyme proteins.

#### Single cell RNA-Seq data processing

We downloaded the raw scRNAseq data and processed it according to Smillie et al^27^. 51 cell types have been characterized by average gene expressions in healthy (n=12) state and non-inflamed UC (n=18). A gene was considered expressed if its log2 expression value was above the mean minus 2 standard deviations of the expressed genes within the cell type.

#### Specific interactions between cell types

We examined all possible connections among the selected 5 cell types. We considered interactions condition specific if they appeared either only in healthy or in UC, i.e. at least one member expressed only in the given condition. We counted the unique PPIs between each cell pair in the two conditions separately (Figure 4c).

#### Cell type specific network of regulatory T cell and downstream pathway analysis

To highlight the downstream effect connections from myofibroblasts to regulatory T cells, we created a cell specific signalling network and we carried out a pathway enrichment analysis. We used the *OmniPath* Cytoscape application^53^ to combine the gene expression data with the *OmniPath* network. We limited the network to genes expressed in regulatory T cells. We selected the receptors targeted by condition-specific ligand-receptor connections in regulatory T cells. Finally, we pruned the network to the two steps neighborhood of the T cell specific receptors. We performed a pathway enrichment analysis on the network described above, using the online interface of the Reactome database with its default settings (hypergeometric test, Benjamini-Hochberg FDR correction, the human genome as the universe).

## Software and data availability

*OmniPath* is available via the Python package *pypath* (https://github.com/saezlab/pypath), the web resource (https://omnipathdb.org), the R/Bioconductor package *OmnipathR* (https://saezlab.github.io/OmnipathR) and the *OmniPath* Cytoscape plug-in^53^. In addition, *pypath* is able to export the network and the enzyme-PTM databases in BEL (Biological Expression Language) format^54^, as well as to generate input files for CellPhoneDB. The BEL format databases are available in BEL Commons^55^. Code is licensed open source (GPLv3). *Pypath* builds the *OmniPath* databases directly from the original resources, hence it gives the highest flexibility for customization and the richest API for queries and manipulation among all access options. Furthermore, it is possible to convert each database to a plain data frame and export in a tabular format. *Pypath* also generates the web resource’s contents which is accessible for any HTTP client at https://omnipathdb.org. Information about the resources is available at https://omnipathdb.org/info. *OmnipathR* and the *OmniPath Cytoscape* plug-in work from the web resource data with convenient post-processing features. All data in *OmniPath* carry the licenses of the original resources (Supplementary Table S12), for profit users can easily limit their queries to fit the legal requirements. A comprehensive guide for *pypath* is available at https://pypath.omnipathdb.org/notebooks/pypath_guide.html.

Apart from the figures presented in this paper, further regularly updated statistics and visualizations are available at https://insights.omnipathdb.org.

A Python and R package for producing the figures and tables of this paper is available at https://github.com/saezlab/omnipath_analysis. The code to build and train the NicheNet method^8^ exclusively using *OmniPath* resources as well as to reproduce the *SARS-CoV-2* case study is freely available at https://github.com/saezlab/NicheNet_Omnipath. The code for building the cell type specific inter- and intracellular networks is available at https://github.com/korcsmarosgroup/uc_intercell.

## Acknowledgements

This work was partially supported by the JRC-COMBINE, partially funded by Bayer, the European Union Innovative Medicines Initiative TransQST (agreement No. 11603), the Federal Ministry of Education and Research (BMBF, Computational Life Sciences grant no. 031L0181B), the DFG (Deutsche Forschungsgemeinschaft / German Research Council; Funding code: SA 3554/1-2, to J.S.R.). T.K. and D.M. were supported by the Earlham Institute (Norwich, UK) in partnership with the Quadram Institute (Norwich, UK) and strategically supported by the UKRI Biotechnological and Biosciences Research Council (BBSRC) UK grants (BB/J004529/1, BB/P016774/1, and BB/CSP17270/1) and by a BBSRC ISP grant for Gut Microbes and Health BB/R012490/1 and its constituent projects, BBS/E/F/000PR10353 and BBS/E/F/000PR10355. L.G was supported by w the BBSRC Norwich Research Park Biosciences Doctoral Training Partnership grant number BB/M011216/1. Thanks to Mirjana Efremova, Daniel Domingo-Fernandez, Charles Tapley Hoyt, Lu Li and Paul D Thomas for their helpful feedback and discussions.

## Authors contributions

D.T. designed and developed *pypath* and *OmniPath* and created the descriptive figures and tables. A.V., D.T. and A.G. developed the *OmnipathR* package. A.V. designed and carried out the case study on *SARS-CoV-2* infection data. L.G. and D.M. designed and carried out the case study on ulcerative colitis. N.P. and O.I. and L.G. contributed to the development of *pypath* and visualization of the database contents. J.S.R. supervised the project, with support from T.K. All authors contributed to the writing of the manuscript.

## Supplementary Materials

**Supplementary Figure S1:**
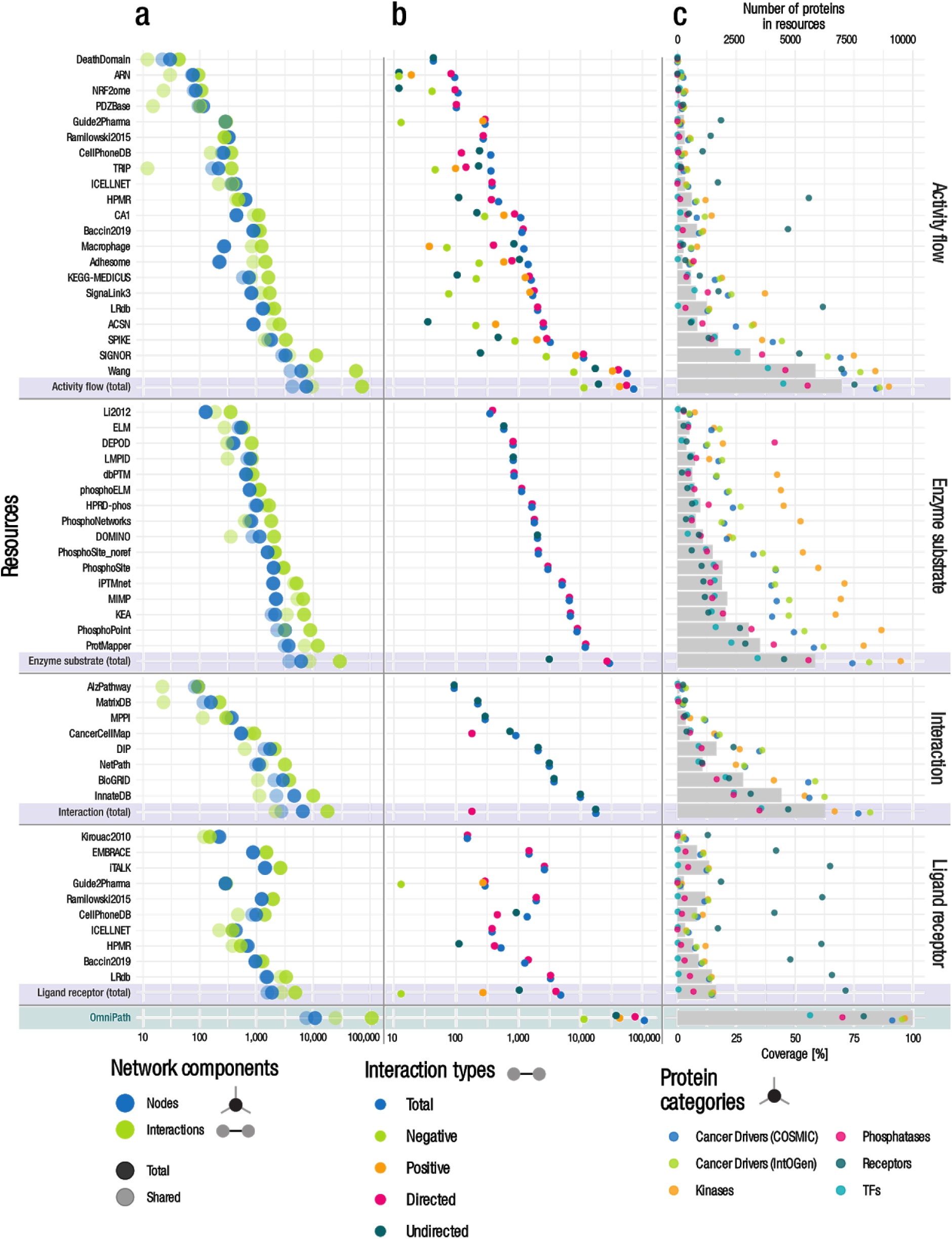
Quantitative description of the PPI network by resource. **(a)** Number of nodes and interactions. The light dots represent the shared nodes and edges (in more than one resource), while the dark ones show their total numbers. **(b)** Causality: number of connections by direction and effect sign. **(c)** Coverage of the networks on various groups of proteins. Dots show the percentage of proteins covered by network resources for the following groups: cancer driver genes from COSMIC and IntOGen, kinases from kinase.com, phosphatases from Phosphatome.net, receptors from the Human Plasma Membrane Receptome (HPMR) and transcription factors from the TF census. Gray bars show the number of proteins in the networks.

**Supplementary Figure S2:**
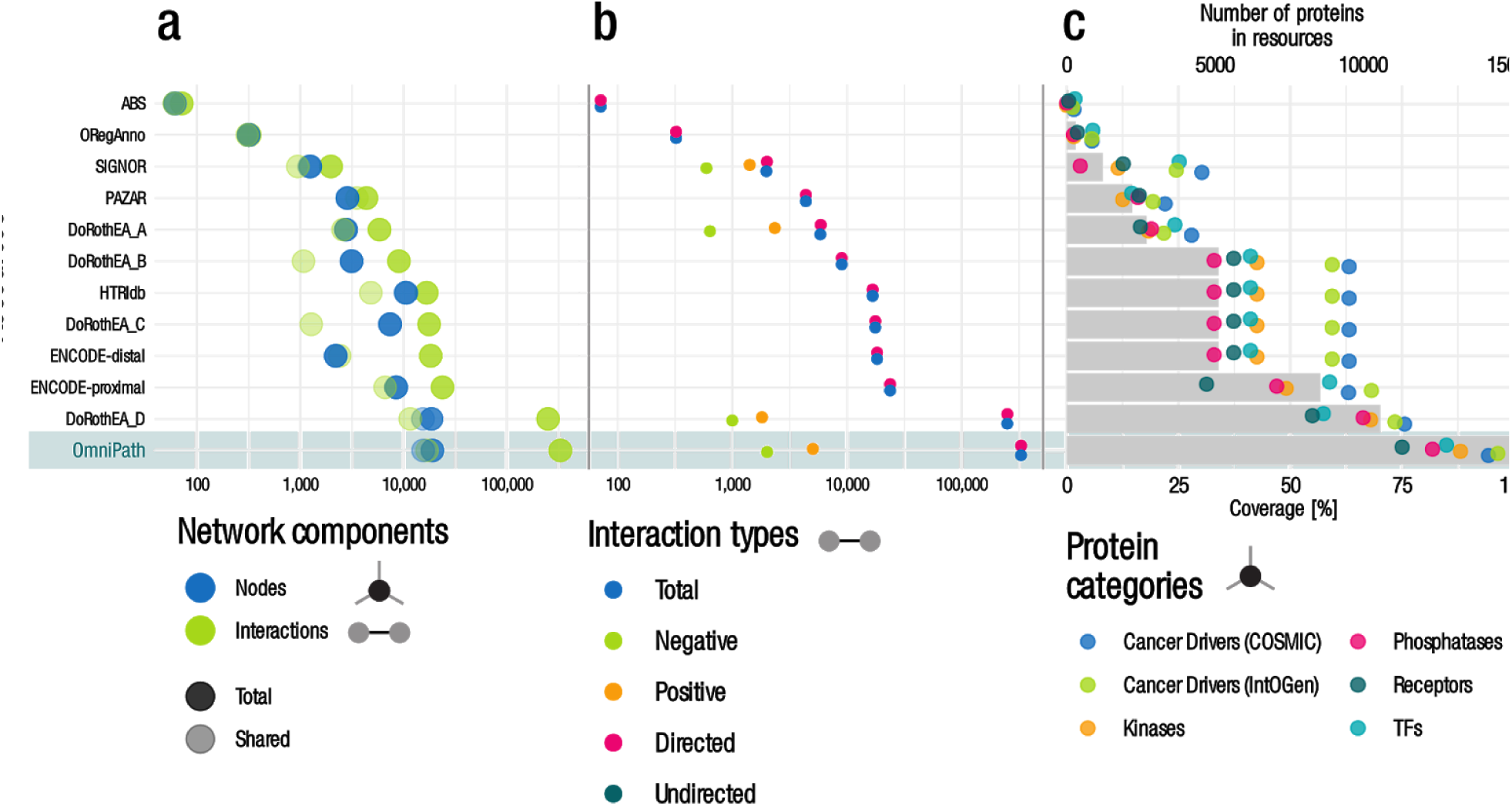
Quantitative description of the transcriptional network by resource. Panels and notations are the same as on Supplementary Figure S1.

**Supplementary Figure S3:**
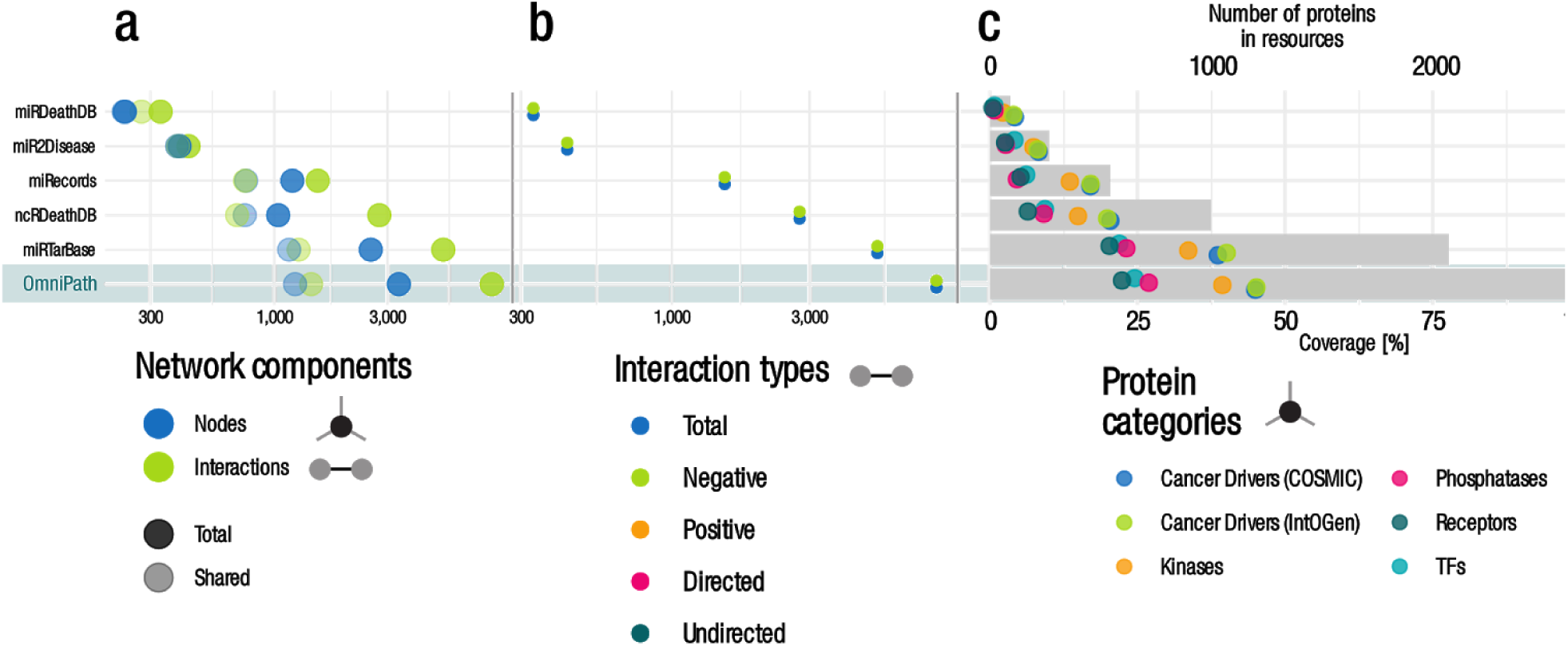
Quantitative description of the post-transcriptional network by resource. Panels and notations are the same as on Supplementary Figure S1.

**Supplementary Figure S4:**
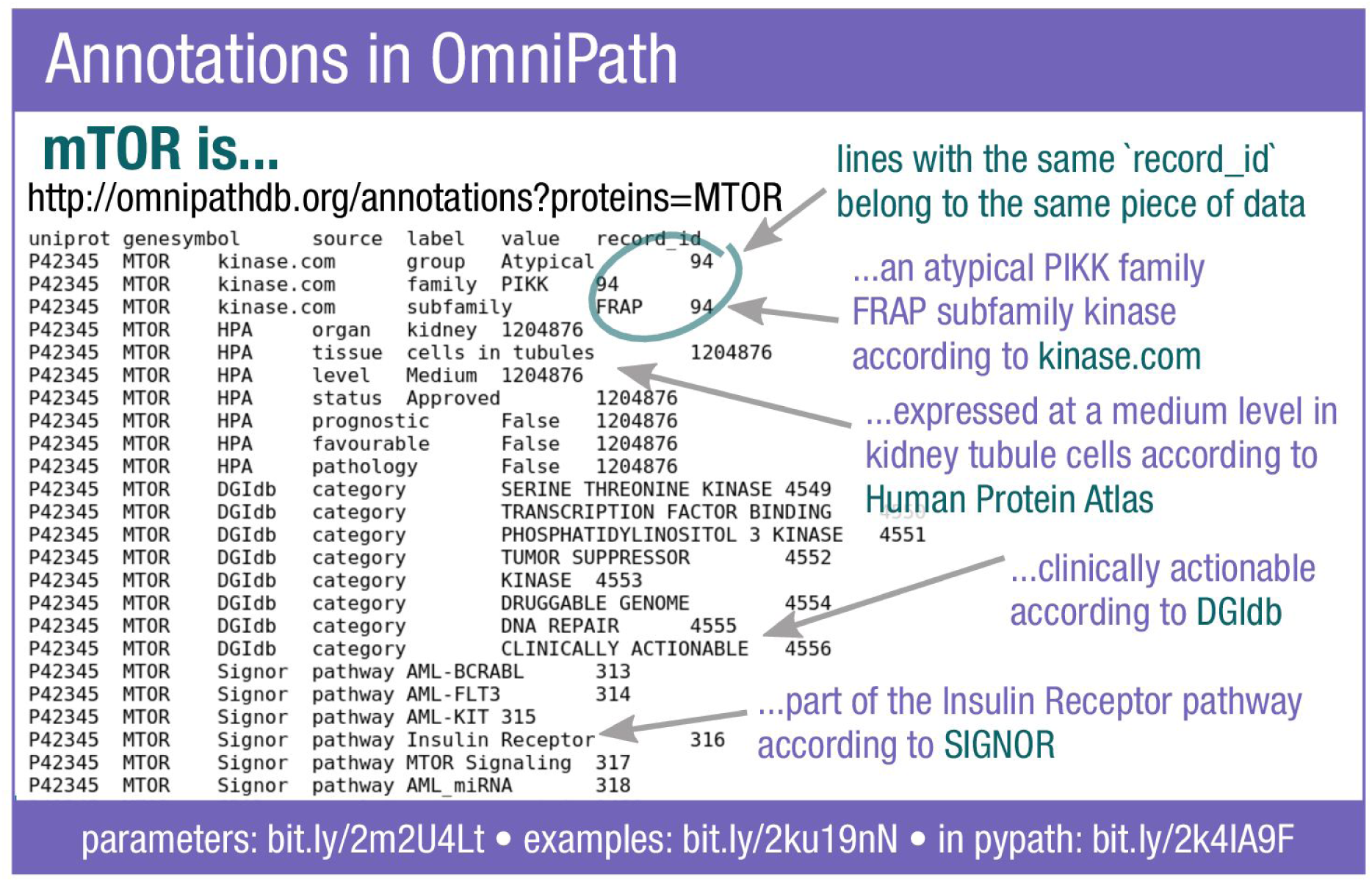
Example of the annotations query in the OmniPath web service. For the protein mTOR a large variety of information is available from different databases. The ‘record_id’ binds together the fields of the record from the original resource. Each field has a ‘label’ and a ‘value’.

**Supplementary Figure S5:**
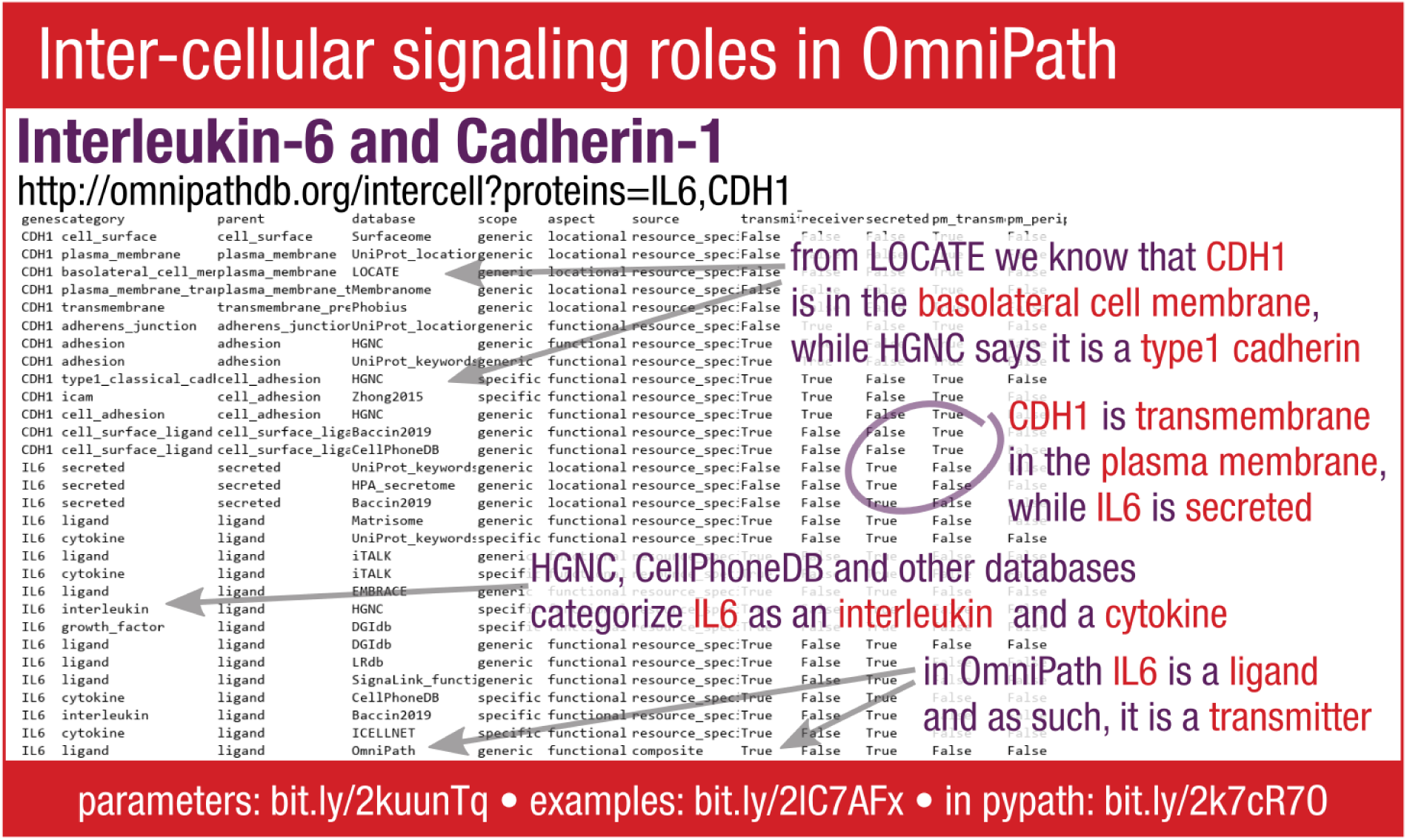
Example of the intercell query in the OmniPath web service. Each category has a parent category and a database of origin. The scope of a category is either ‘generic’ (e.g. ligand) or ‘specific’ (e.g. interleukin). The aspect is either ‘locational’ or ‘functional’. Further attributes show whether the protein is a signal transmitter or a receiver, and whether it is secreted, or a transmembrane or peripheral protein of the plasma membrane.

**Supplementary Figure S6:**
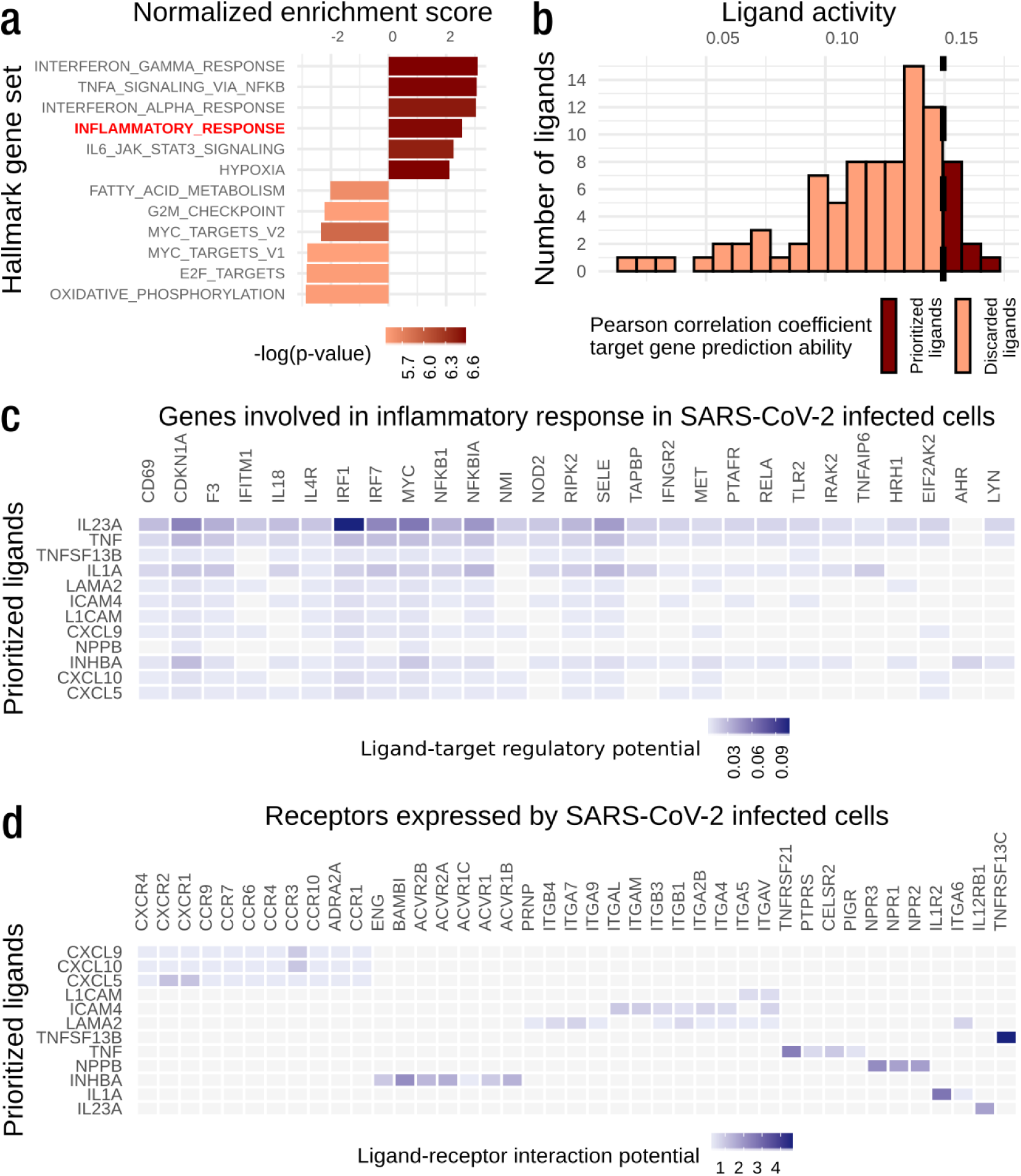
OmniPath-based NicheNet analysis to predict overexpressed ligands in SARS-CoV-2 infection potentially affecting the expression of inflammatory response related genes in Calu3 cells. (a) Most significantly enriched gene sets after SARS-CoV-2 infection on the Calu3 cell line. Inflammatory response is highlighted in red. (b) Results of NicheNet’s ligand activity analysis: Number of overexpressed ligands after SARS-CoV-2 infection and their potential to predict the inflammatory response gene set based on the Pearson correlation coefficient. The top 12 ranked ligands, out of a total of 117 overexpressed ligands, were selected. (c) Regulatory potential of the top ranked ligands and target genes from the inflammatory response program based on NicheNet’s prior knowledge model. (d) Ligand-receptor interaction potential based on NicheNet’s prior knowledge model between the top ranked ligands and the receptors expressed in the Calu3 cell line.

***Supplementary Table S1: List of resources in OmniPath and pypath*.**

*Besides the name, webpage and publication of the resources we list which ones of the five major OmniPath databases (network, enzyme-PTM, complexes, annotations, intercell) each resource contributes to, and which datasets within the network database. Certain resources are not redistributed by the OmniPath web service or not used for any of the databases, but available only by pypath or used for different purposes such as identifier translation, curation facilitation, etc*.

***Supplementary Table S2: Quantitative description of the OmniPath network database.***

*Number of shared (overlap with any other resource) and unique molecular entities in total and by entity type, number of interactions in total, and by direction and effect sign, number of references and curation effort (unique reference-interaction pairs). Total rows are shown for each dataset and interaction type (PPI, transcriptional, post-transcriptional, TF-miRNA). In the total rows the components are counted as shared if they can be found in more than one resource*.

***Supplementary Table S3: Quantitative description of the OmniPath enzyme-PTM database.***

*Number of shared (overlap with any other resource) and unique enzyme-PTM relationships, references and curation effort (reference-record pairs), list of available modification types*.

***Supplementary Table S4: Quantitative description of the OmniPath complexes database.***

*Number of protein complexes, homomers and heteromers, shared (overlap with any other resource) and unique records, availability of stoichiometry information, number of references and curation effort (reference-record pairs)*.

***Supplementary Table S5: Quantitative description of the OmniPath annotations database.***

*Each record carries the attributes listed in the ‛fields‛ column. At resources where no attributes are listed here, the annotation can be considered a set, i.e. a molecular entity either belongs to this set or not. One molecular entity might have more than one annotation records from the same resource*.

***Supplementary Table S6: Quantitative description of the OmniPath intercell database.***

*Size and contents of the generic functional and locational categories in the intercellular communication roles (intercell) database. Functional categories are either transmitters, receivers or both; locational don’t have these attributes. ‛OmniPath’ in the resources column appears if certain subclasses of the category are defined directly by OmniPath not by an integrated resource. The specific categories are not shown in this table*.

***Supplementary Table S7: Secondary resources in OmniPath.***

*Some resources integrate data from other resources. In OmniPath the records carry information both about the primary (directly integrated into OmniPath) and the secondary resources*.

***Supplementary Table S8: Size of the transcriptional regulatory network.***

*Number of nodes, interactions, transcription factors and target genes are shown for networks of interactions with or without literature references, using DoRothEA confidence levels A-D vs. A-E. All networks include, apart from DoRothEA, other resources integrated directly into OmniPath: ABS, ENCODE, HTRI, ORegAnnO, PAZAR, SIGNOR*.

***Supplementary Table S9: Success rate of homology translation.***

*Here we show the success rate of homology translation of the OmniPath human PPI signaling network to mouse and rat using the NCBI HomoloGene database. Number and percentage of nodes (genes) and interactions successfully translated*.

***Supplementary Table S10: Terminology in the intercellular communication roles (intercell) database.***

*The attributes category name, parent category, source, aspect and scope are carried by each category in the intercell database. Below we define the possible values of these attributes. We also define here those major categories (e.g. secreted, receptor, etc) which are pivotal for an unambiguous definition of the intercellular communication roles*.

***Supplementary Table S11: Dominant pathways in healthy and ulcerative colitis networks.***

*The networks have been created from condition specific receptors and proteins within two steps from the receptors. Using the Reactome database, we highlighted the top ten pathways*.

***Supplementary Table S12: Licensing terms of the resources in OmniPath.***

*The license field is highlighted in green if the resource is freely available for commercial (for-profit) use, in yellow if only for academic or non-profit use, and in grey if we are awaiting clarification from the copyright holders. In the OmniPath interfaces (pypath, web service, OmnipathR) users are able to set their license preferences to ensure their data usage meets the legal requirements*.

## Supplementary Note 1: Joint analysis of intra- and intercellular processes in SARS-CoV-2 infected lung epithelial cancer (Calu3) cells

In this note, we provide further details and supporting literature for the results obtained in the *SARS-CoV-2* case study and presented in Figure 4b and Supplementary Figure S6. In this case study, we aim to explore the potential autocrine regulatory effect of ligands overexpressed in *SARS-CoV-2* infection of epithelial lung cancer cells *(Calu3)* on the expression of inflammatory response genes. We used expression data from a recent publication^1^.

We first performed a differential expression analysis of *SARS-CoV-2* infected cells versus mock treated controls. This allowed us to carry out a gene set enrichment analysis revealing inflammatory response as one of the most enriched sets (Supplementary Figure S6a). We subsequently selected the most relevant genes involved in inflammatory response (Methods). In addition, we selected over-expressed ligands after infection that are likely to be secreted to the extracellular milieu (Methods). We then applied our *OmniPath*-based version of *NicheNet* to rank the overexpressed ligands secreted by infected *Calu3* cells that are most likely to be involved in the regulation of inflammatory response related genes (Methods). Out of a total of 117 overexpressed ligands, we selected the 12 top-ranked ones for subsequent analysis according to the distribution of correlation values (Supplementary Figure S6b) and *nichenetr* guidelines^2^. Among them, we found different types of cytokines: interleukins (*IL23A* and *IL1A*), tumor necrosis factors (*TNF* and *TNFSF13B*) and chemokines (*CXCL5*, *CXCL9* and *CXCL10*). These proteins are known to be involved in the immune and inflammatory response, hence supporting our *OmniPath*-based approach. Indeed, we evaluated to which extent our top 12 prioritized ligands can together predict whether the top predicted targets belong to our previously defined inflammatory response gene set or not (average p-value=3.25e-08 from Fisher’s exact tests after 10 cross-validation rounds). We can therefore assume that the overexpressed ligands secreted after *SARS-CoV-2* infection can explain, at least to some extent, the expression of inflammatory response related genes in the *Calu3* cells.

*NicheNet* ranks the ligands based on their potential effect to regulate the whole set of inflammatory response genes^2^. In order to get more detailed functional and mechanistic insights, we next investigated the inter- and intracellular signaling events that can lead to the activation of a particular ligand-target link. First, we explored the *NicheNet* regulatory potential scores between our top-scored ligands and the top inflammatory response target genes according to our *OmniPath*-based prior knowledge network (Supplementary Figure S6c). Then, we selected the receptors expressed in *Calu3* cells after infection that can potentially bind our top ranked ligands, i.e. a known interaction is described between them in our ligand-receptor network (methods). The most likely ligand-receptor pairs according to their *NicheNet* prior interaction potential score are displayed in Supplementary Figure S6d. We finally inferred the most likely paths connecting some of our top ranked ligands to their inflammatory response target genes (Figure 4b and methods).

Among the top predicted ligands, we found three C-X-C motif chemokines (*CXCL5*, *CXCL9* and *CXCL10*). *CXCL9* and *CXCL10* are well known pro-inflammatory chemokines that participate in the inflammatory response by recruiting immune cells to infected areas^3^. According to our results, these ligands may potentially bind to C-X-C chemokine receptors (*CXCR1* and *CXCR2*) and to the *CCR3* receptor (Figure 4b). Then, *CXCR1* and *CCR3* can both activate *MAPK14*, a serine/threonine kinase which plays a key role in the signalling responses to extracellular stimuli such as proinflammatory cytokines or physical stress leading to direct activation of transcription factors^4^. In addition, *CXCR1, CXCR2* and *CCR3* directly interact with *JAK2*, activating the STAT transcription factors. In particular, *JAK2* mediates the cytokine-driven activation of the *FOS* transcription factor, which is a key component in the regulation of proinflammatory genes^5^. Consequently, the use of ruxolitinib, a *JAK1* and *JAK2* inhibitor, has been suggested as a potential way to prevent the harmful effects of the excessive secretion of proinflammatory proteins, the so-called *cytokine storm*, in severe cases of *COVID-19*^6^.

We also identified two interleukins (*IL23A* and *IL1A*) among the top predicted ligands. *IL23A* forms a heterodimeric cytokine by associating with IL12B, the IL-23 interleukin. IL-23 binds to the *IL12RB1*-*IL23R* receptor complex and activates the JAK-STAT signaling cascade promoting the production of proinflammatory cytokines. Furthermore, IL-23 induces autoimmune inflammation and its inhibition is the main treatment for *psoriasis*, an autoimmune disease^7^. In our results (Figure 4b), we identified the interaction between *IL23A* and *IL12RB1*, and how *IL12RB1* directly activates some of the STAT transcription factors (*STAT1, STAT3* and *STAT4*). *IL1A* is known to play key roles in the regulation of the immune and the inflammatory response. It binds to the interleukin-1 receptor, interaction that was partially recovered in our signaling network (*IL1R2*, Figure 4b). Then, *IL1R2* activates *CASP3*, whose role in the modulation of cytokine expression and inflammation has been proposed^8^, although is not as straightforward as in the previous discussed examples.

Finally, we also retrieved some tumor necrosis factors (*TNF* and *TNFSF13B*) as top ligands potentially regulating the expression of inflammatory response related genes. The main functions of *TNF* are the regulation of immune cells and the systemic inflammatory response. Once *TNF* comes to contact with their potential receptors, the *TRADD* protein can also bind to the receptor resulting in the potential initiation of three different pathways: activation of the NFKB pathway, activation of the MAPK pathway or induction of death signaling^9^. Our results capture the interaction between *TNF* and *TRADD* to their potential receptor, *TNFRSF21*, which in turn activates *RELA* (Figure 4b). The activation of *RELA* suggests an activation of the *NF-kB* pathway, known to be active in *SARS-CoV-2* infection^10^. The *TNFSF13B* gene encodes the B-cell activating factor (*BAFF*) protein, which can bind to the *TNFRSF13C* receptor as identified in our results. The interaction between *BAFF* and its receptors triggers the activation of the classical and non-canonical *NF-kB* signaling pathway^11^. In our results, we identified the activation of *MAP3K14*, which indeed appears to be involved in the activation of the *NF-kB* complex and its transcriptional activity^12^.

In summary, we studied how the ligands secreted after SARS-CoV-2 infection could influence the inflammatory response of neighboring cells. We were able to capture known biological processes supported by the literature. These processes and signaling cascades may lead to the exacerbated inflammatory response observed in COVID-19 most severe cases.

